# Chemerin-CMKLR1 mediated OGD/R induced Mitochondrial Dysfunction, Oxidative Stress, and Autophagy differentially in Microglia and Neurons

**DOI:** 10.1101/2025.07.28.667159

**Authors:** Pei-Yan Long, Zhi Tang, Na Cai, Zhi-Qin Yan, Xiao Gao, Zheng-Wei Wang, Zhi-Zhong Guan, Xiao-Lan Qi, Ruiqing Ni, Yan Xiao

**Affiliations:** Key Laboratory of Endemic and Ethnic Diseases, Ministry of Education & Key Laboratory of Medical Molecular Biology of Guizhou Province, Guizhou Medical University, Guiyang, Guizhou Province, China; Gui Qian International General Hospital Guiyang, Guizhou Province, China; Collaborative Innovation Center for Prevention and Control of Endemic and Ethnic, Guiyang, Guizhou Province, China; Institute for Regenerative Medicine, University of Zurich, Zurich, & Department of Nuclear Medicine, Inselspital, Bern, Switzerland

**Keywords:** apoptosis, autophagy, chemerin, CMKLR1, ischemic stroke, microglia, mitochondria, neuron, oxidative stress, oxygen and glucose deprivation/reoxygenation (OGD/R)

## Abstract

**Introduction:** Ischemia-reperfusion (I/R) injury exacerbates tissue damage upon reperfusion after ischemia. The impact of chemerin and its receptor, chemokine-like receptor 1 (CMKLR1) on I/R injury remains poorly understood. We hypothesized that chemerin-CMKLR1 differentially regulates signaling in microglia and neuronal cells oxygen-glucose deprivation/reoxygenation (OGD/R), influencing mitochondrial function, oxidative stress, and autophagy.

**Methods:** Using BV2 microglia and Neuro-2a (N2a) neuronal cells, we examined OGD/R-induced changes in autophagy associated proteins, chemerin and CMKLR1 expression. We investigated the functional consequences of CMKLR1 overexpression and chemerin treatment on oxidative stress, apoptosis, autophagy, and mitochondrial dynamics in BV2 microglia and N2a neuronal cells.

**Results:** OGD/R downregulated CMKLR1 while upregulating autophagy in both BV2 microglia and N2a cells; While chemerin expression decreased in BV2 microglia but increased in N2a cells following OGD/R. Treatment with chemerin dose-dependently reduced oxidative stress and apoptosis while enhancing mitochondrial fusion, suppressing fission, and promoting autophagy and mitochondrial function in both cell types under OGD/R. CMKLR1 overexpression exacerbated mitochondrial respiratory dysfunction, mitochondrial fusion, fission, and elevated autophagy (LC3II/LC3I and Pink1 levels), with cell-type-specific differences observed in Parkin and P62 regulation.

**Conclusion:** Our study demonstrates cell-type-specific regulation of chemerin-CMKLR1 signaling in I/R injury, and distinct mitophagy activation mechanisms in microglia and neurons. These findings suggest cell-type specific modulation of chemerin-CMKLR1 as a potential therapeutic target in preserving mitochondrial homeostasis, modulating autophagy, mitophagy and reducing oxidative stress, apoptosis in both microglia and neurons for mitigating I/R injury.

## INTRODUCTION

Stroke is the second leading cause of death and a significant contributor to disability worldwide [1]. Ischemia/reperfusion (I/R) injury is caused by partial or complete cessation of blood flow after blood vessel infarction, resulting in a significant decrease in available oxygen and energy substrates and sudden restoration of blood flow to prevent further ischemia [2]. I/R injury may lead to further cell and tissue damage and impair the functional outcome of patients with recanalization after thrombectomy [3]. Timely restoration of blood flow and oxygen after obstruction is extremely critical for maintaining the normal structure and function of nerve cells. I/R injury is associated with multiple pathological processes, including inflammatory responses, oxidative stress, endoplasmic reticulum stress, mitochondrial dysfunction and apoptosis [4, 5]. Microglia is the resident immune cell and exert time-dependent dual roles in ischemic stroke. After injury, anti-inflammatory microglia promote recovery through neurotrophic factor release, whereas later pro-inflammatory polarization exacerbates damage via inflammatory mediators [6]. Microglia-mediated crosstalk with neuron and other cells in the brain and play pivotal role for functional recovery post-stroke. The understanding of the mechanisms underlying I/R injury is still limited, which hinders the development of effective therapeutic interventions [7, 8].

Chemerin is an inflammatory factor or chemokine that is mainly secreted by adipose tissue and is synthesized and secreted by liver, kidney and placental tissue [9–11]. Chemokine-like receptor 1 (CMKLR1 in animals and ChemR23 in humans) is a G protein-coupled receptor and one of the three receptors for chemerin [12]. The chemerin-CMKLR1 axis is involved in multiple intracellular signaling pathways and downstream inflammatory factors, mitophagy, angiogenesis, glycolipid metabolism, and apoptosis [13–17]. Chemerin regulates local and systemic inflammation and induces immune cell chemotaxis, which can promote the migration and aggregation of immune cells, thereby leading to the secretion of a variety of inflammatory factors, such as interleukin-6, tumor necrosis factor-α and C-reactive protein, to aggravate the inflammatory response [18–20]. Chemerin has a complex role in neuroinflammation, with evidence suggesting that it can either increase or decrease neuroinflammation. Earlier studies have demonstrated that preischemic treatment with recombinant human chemerin has a neuroprotective effect on rat and mouse models of ischemic stroke and hypoxic-ischemic encephalopathy by reducing oxidative stress and apoptosis [21–23]. Synthetic chemerin-derived peptides can also suppress inflammation through ChemR23 [24]. In addition, chemerin can promote mitophagy [25], autophagy and endoplasmic reticulum stress [26]. The chemerin–CMKLR1 axis is functionally important for the central regulation of energy homeostasis [27] and is involved in I/R injury [14]. Studies have shown that chemerin is expressed in the serum of patients with ischemic stroke [28]. It also stimulates aortic smooth muscle cell proliferation and migration via the activation of autophagy in metabolic hypertension rats [25]. A reduction in CMKLR1 has been shown to reverse the mitochondrial dysfunction caused by chemerin [29]. ChemR23 signaling has been shown to ameliorate brain injury via the inhibition of oxidative stress and NLRP3 inflammasome-mediated neuronal pyroptosis in ischemic stroke [30] and cognitive impairments in diabetic mice [31]. The chemerin-ChemR23 signaling axis is involved in endothelial protection via the adenosine triphosphate (ATP)-sensitive potassium channel opener iptakalim [32]. CMKLR1 deficiency influences glucose tolerance and thermogenesis in high-fat diet-fed mice [14, 33].

The role of chemerin and its receptor CMKLR1 in I/R injury is unclear. Oxygen and glucose deprivation reoxygenation (OGD/R) is a commonly used in vitro model that mimics I/R injury in the brain. We hypothesized that OGD/R can induce cell type-specific effects via chemerin-CMKLR1 signaling in BV2 microglia and N2a cells and lead to functional consequence. OGD/R has been shown to induce mitochondrial dysfunction and oxidative stress in neurons [34] and proinflammatory, oxidative, and metabolic responses in microglia [35]. Here, we further investigated the effect of chemerin treatment and CMKLR1 overexpression on oxidative stress, mitochondrial function, autophagy and apoptosis in BV2 microglia and N2a cells.

## 2. MATERIALS AND METHODS

### 2.1 Materials and Antibodies

Mouse microglia (BV2 cells) were donated by Professor Wenfeng Yu’s research group at the Key Laboratory of Molecular Biology of Guizhou Medical University, and mouse-derived neuroblastoma N2a cells were donated by Professor Jun Tan’s research group at the Key Laboratory of Molecular Biology of Guizhou Medical University. The lists and sources of materials, including antibodies, chemicals and kits, are described in detail in **STable 1** and **STable 2**. Mouse recombinant chemerin protein (20 μg) was dissolved in sterile water, further diluted with 0.1% bovine serum albumin, aliquoted, stored in a -80°C freezer, and prepared with complete medium to the desired concentration.

### 2.2 Cell culture and determination of the time of OGD/R

BV2 microglia and N2a cells were cultured in 89% (v/v) Dulbecco’s modified Eagle medium (DMEM) supplemented with 10% (v/v) fetal bovine serum and 1% (v/v) penicillin‒streptomycin at 37°C. At 80%-90% confluence, the cells were seeded into 6-well culture plates at a density of 1.5×10^5^ cells/well and incubated at 37°C with 5% CO_2_ for 24 h. The medium of the control group was replaced with normal medium, 3 h/24 h group (oxygen and glucose deprivation for 3 h and then reoxygenation culture for 24 h) and 6 h/24 h group (oxygen and glucose deprivation for 6 h and then reoxygenation culture for 24 h) with 1% penicillin‒streptomycin glucose-free DMEM in a hypoxic chamber with 1% O_2_, 94% N_2_ and 5% CO_2_ and at 37°C for 3 h or 6 h. Finally, the medium was discarded, and the cells were cultured for an additional 24 h for reperfusion in normoxia, glucose and fetal bovine serum-containing media under 37°C, 5% CO_2_ conditions.

### 2.3 Cell viability experiments

The cells were seeded in 96-well plates at 5 ×10^3^ cells/well in 100 μL of culture medium. When the cells reached a density of 70%, they were deprived of oxygen and glucose for 6 h and then treated with various concentrations of mouse recombinant chemerin protein (10, 20, 40, 60, 80, 100 and 120 ng/mL) for 24 h at 37°C and 5% CO_2,_ as described previously [36]. The medium was discarded, 10 μL of Cell counting kit-8 reagent was mixed with 100 μL of complete medium in each well, and the cells were incubated at 37°C for 1 h. The absorbance was measured at 450 nm via a microplate reader (Bio-Rad, Hercules, USA).

### 2.4 OGD/R model and chemerin treatment

The cells were seeded into 6-well culture plates and incubated at 37°C and 5% CO_2_ for 24 h. The medium of the control group was replaced with complete medium. The media for the OGD/R and OGD/R+chemerin groups were replaced with 1% penicillin‒streptomycin glucose-free DMEM in an incubator at 37°C, 1% O_2_, 94% N_2_ and 5% CO_2_ for 6 h. After this, the media for the OGD/R+chemerin group were replaced with complete media, and those for the OGD/R+chemerin group were replaced with complete media containing recombinant chemerin protein at concentrations of 40 ng/mL, 100 ng/mL and 120 ng/mL. The cells were collected after 24 h of culture at 37°C and 5% CO_2_.

### 2.5 Lentivirus transfection and OGD/R model

BV2 microglia and N2a cells were transfected with the green fluorescent protein-containing puromycin-resistant recombinant lentivirus CMKLR1 and empty lentiviral vectors at a multiplicity-of-infection of 100 (Genechem, Shanghai, China). Three days later, culture medium containing 5 μg/mL puromycin was added to the cell culture plates. The efficiency of CMKLR1 overexpression was estimated by real-time quantitative PCR (qPCR) and Western blotting after seven days. The infected cells were subsequently cultured in media supplemented with 2.5 μg/mL puromycin. Cells stably overexpressing CMKLR1 were cultured in normal complete medium for 24 h and then cultured with sugar-free DMEM. The cells were placed in a hypoxic chamber for 6 h. Finally, the medium was discarded, and the cells were cultured for an additional 24 h for reperfusion in oxygen, glucose and fetal bovine serum-containing media under normal conditions.

### 2.6 Enzyme-linked immunosorbent assay

To measure the concentration of chemerin in the cell culture, we collected the cell culture medium supernatant and measured it using a sandwich enzyme-linked immunosorbent assay (ELISA) for mouse chemerin (4A Biotech, Beijing, China, #CME0119) as described previously. The absorbance was measured at 450 nm using a microplate reader (Bio-Rad, Hercules, USA).

### 2.7 Western blotting analysis

Cells (n=3 cell samples in each group) were treated, homogenized and lysed in ice-cold radioimmunoprecipitation assay lysis buffer supplemented with 1 mM phenylmethanesulfonyl fluoride. Protein concentrations were measured via a BCA protein assay kit (Takara, Japan, #T9310A). Equal amounts (20–40 μg) of protein were separated via 10–15% sodium dodecyl sulfate‒polyacrylamide gel electrophoresis, and then blotted onto polyvinylidene difluoride membranes (Millipore, MA, USA; #ISEQ00010). Next, the membranes were blocked with 5% skim milk in Tris-buffered saline with Tween for 1 h at room temperature and further incubated at 4°C overnight with the following primary antibodies: CMKLR1, PTEN-induced kinase 1 (Pink1), Parkin, dynamin-related protein 1 (Drp1), fission 1 protein (Fis1), mitofusin-1 (Mfn1), mitofusin-2 (Mfn2), sequestosome-1 (P62), and β-actin, as described previously [37] (list of antibodies in **STable 1**). After three washes with Tris-buffered saline with Tween, the membranes were incubated with mouse and rabbit secondary antibodies at room temperature for 1 h. The immunoreactive bands were detected via enhanced chemiluminescence. The signal intensity was quantified via ImageJ software (NIH). The signal was normalized to that of β-actin.

### 2.8 Immunofluorescence Staining

N2a cells and BV2 microglia were grown on coverslips in 12-well or 24-well plates. The cells were fixed with 4% paraformaldehyde for 20 min. After permeabilization with 1× phosphate-buffered saline (containing 0.2% Triton X-100) for 10 min, the cells were blocked with goat serum for 30 min. The coverslips were incubated with primary antibodies against 4-hydroxynonenal (4-HNE) and 8-hydroxydeoxyguanosine (8-OHdG) overnight at 4°C in the dark, as described previously [38]. Coverslips with goat anti-mouse IgG (H+L) Cyanine3 and goat anti-rabbit IgG (H+L) Alexa Fluor™ 488 were incubated for 1 h at room temperature in the dark. Nuclei were counterstained with 4′,6-diamidino-2-phenylindole. Images were acquired by using laser confocal microscopy (Olympus SpinSR10, Japan). The relative fluorescence intensity of 50–70 cells in each group was analyzed by using ImageJ.

### 2.9 Reactive oxygen species (ROS) measurements

N2a and BV2 cells were collected in 1.5 mL eppendorf tubes by ethylenediaminetetraacetic acid-free trypsinization and centrifuged at 1000 rpm/min for 5 min. Dihydroethidium staining solution from the ROS assay kit was diluted with DMEM (1:1000 v/v, final concentration of 5 μM). A total of 500 μL of dihydroethidium staining solution was added to each tube. The cells were incubated at 37°C for 30 min in the dark as described previously [39]. The staining solution was removed, and the cells were gently washed twice with DMEM. The cells were resuspended in 500 μL of 1× phosphate-buffered saline, and the level of ROS was detected using a flow cytometer (BD Biosciences, USA).

### 2.10 Real-time quantitative PCR analysis

Total RNA was isolated from BV2 microglia and N2a cells using TRIzol reagent according to the standard protocol. Total RNA was reverse-transcribed in 20 μL of a reaction mixture containing RNA, 2× RT-Reaction mix, Starscript II RT mix and diethyl pyrocarbonate -ddH_2_O, using StarScript II RT Mix with gDNA Remover (Genestar, Japan, #A224-10). The assay was run at 42°C for 20 min, 85°C for 5 min, and then stored at 4°C. The quantification of gene expression was performed using real-time qPCR System (Bio-Rad, USA) with 2×RealStar Fast SYBR qPCR Mix (Genestar, Japan, #A301) and specific primers (**STable 3**). The 10 μL total PCR mixture consisted of specific primers (0.25 μM final concentrations), 1 μL of cDNA template, 5 μL of 2 × SYBR Green master mix and ddH_2_O. Amplification was performed under the following cycle conditions: 2 min at 95°C, 40 cycles of 15 s denaturation at 95°C, annealing at 60°C for 30 s and extension at 72°C for 30 s. The threshold cycle (CT) value and the relative mRNA expression of the genes were calculated using the 2^−ΔΔCT^ method.

### 2.11 Mitochondrial DNA (mtDNA) copy numbers

Total DNA was extracted from the cells using a TIANamp Genomic DNA Kit (TIANGEN, China, #DP304-02). Absolute quantitative PCR was performed as previously described [40]. The ratio of mtDNA to nuclear DNA was used to assess the relative mtDNA copy number. The abundances of mitochondrially encoded NADH dehydrogenase 1 (mt-ND1)/β-globin and mitochondrially encoded 12S RNA (mt-RNR1)/β-actin were determined. The sequences of the primers are described in STable 3.

### 2.12 Measurement of mitochondrial bioenergetics

The mitochondrial oxygen consumption rate (OCR) of cells after OGD/R treatment was measured using a Seahorse XFp Analyzer with a Seahorse XF cell mito-stress test kit (Agilent, USA, #103010-100). Briefly, the cells were plated on Seahorse XFp microplates. After 24 h, the cells were subjected to OGD/R and then washed twice with modified DMEM (pH 7.4) supplemented with 2 mM glutamine, 1 mM pyruvate and 10 mM glucose. The assay medium was preequilibrated in an incubator at 37°C without CO_2_ for 1 h, and the cells were incubated in an incubator without CO_2_ at 37°C for 60 min. The microplates were loaded into the analyzer, and three compounds, oligomycin (1.5 μM), carbonylcyanide-4-(trifluoromethoxy) phenylhydrazone (FCCP, 2 μM) and a mixture of antimycin and rotenone A (0.5 μM), were sequentially injected into the samples during the OCR measurement. The OCR values in each well were normalized to the protein concentration.

### 2.13 Mitochondrial ROS detection

The production of mitochondrial ROS in cells after OGD/R treatment was assessed with the MitoSOX red mitochondrial superoxide indicator (Invitrogen, USA, #M36008). The cells were harvested, washed twice with Hank’s balanced salt solution (HBSS), and then incubated in 500 μL of HBSS containing 5 μM MitoSOX for 30 min at 37°C in an incubator. After being washed two times in HBSS, the cells were analyzed using flow cytometry (BD Biosciences, USA).

### 2.14 Detection of apoptosis

The apoptosis of BV2 microglia and N2a cells treated with chemerin was evaluated using annexin V-fluorescein isothiocyanate (FITC)/propidium iodide (PI) and annexin V-allophycocyanin (APC)/7-aminoactinomycin D (7-AAD) apoptosis detection kits (BD Biosciences, USA, #556547; MULTISCIENCES, China, #AP105). Briefly, the cells were co-incubated with 5 μL of Annexin V-FITC and 5 μL of propidium iodide for 15 min at room temperature in a light-free environment and then detected using flow cytometry (BD Biosciences, USA). After OGD/R, the cells were harvested and stained with 5 μL of annexin-V-APC and 10 μL of 7-AAD for 5 min at room temperature in the dark. The percentage of apoptotic cells was analyzed using flow cytometry (BD Biosciences, USA).

### 2.15 Statistical analysis

Welch’s t test was used for two-group comparisons. One-way ANOVA and least significant difference (LSD) post hoc analysis were used for multiple-group comparisons using SPSS 22. The data are presented as the mean±standard deviations (SD) (n=3–5 repeats). Significance was set at *P* < 0.05.

## RESULTS

### 3.1 OGD/R differentially altered chemerin expression in BV2 microglia and N2a cells, while similarly increased autophagy associated protein and reduced CMKLR1 levels

Given the dynamic, time-dependent role microglia in I/R injury, we first established the optimal OGD/R duration (3 h or 6 h OGD) by assessing autophagy markers in BV2 microglia and N2a cells. Autophagy was evaluated by the expression level of P62 and the LC3II/LC3I ratio, which are established indicators of autophagic activity [41]. In BV2 cells, 3 h OGD/24 h reperfusion increased LC3II/LC3I by 33% (*P* = 0.004) and reduced the level of P62 by 62% (*P* = 0.001), while 6 h OGD/24 h reperfusion increased LC3II/LC3I ratio by 81% (*P* = 0.0001) and decreased the level of P62 by 75% (*P* = 0.0002) (**Fig. 1A– C**). Similarly, N2a cells exhibited a more pronounced response with 6 h OGD. LC3II/LC3I ratio was increased by 367% (*P* = 0.008) after 3 h OGD and 1131% (*P* < 0.0001) after 6 h OGD, with P62 level decreased by 15% (*P* = 0.011) and 20% (*P* = 0.003), respectively (**Fig. 1D–F**). Based on these results, the 6 h OGD/24 h reperfusion condition was selected for subsequent experiments.

**Fig 1.**
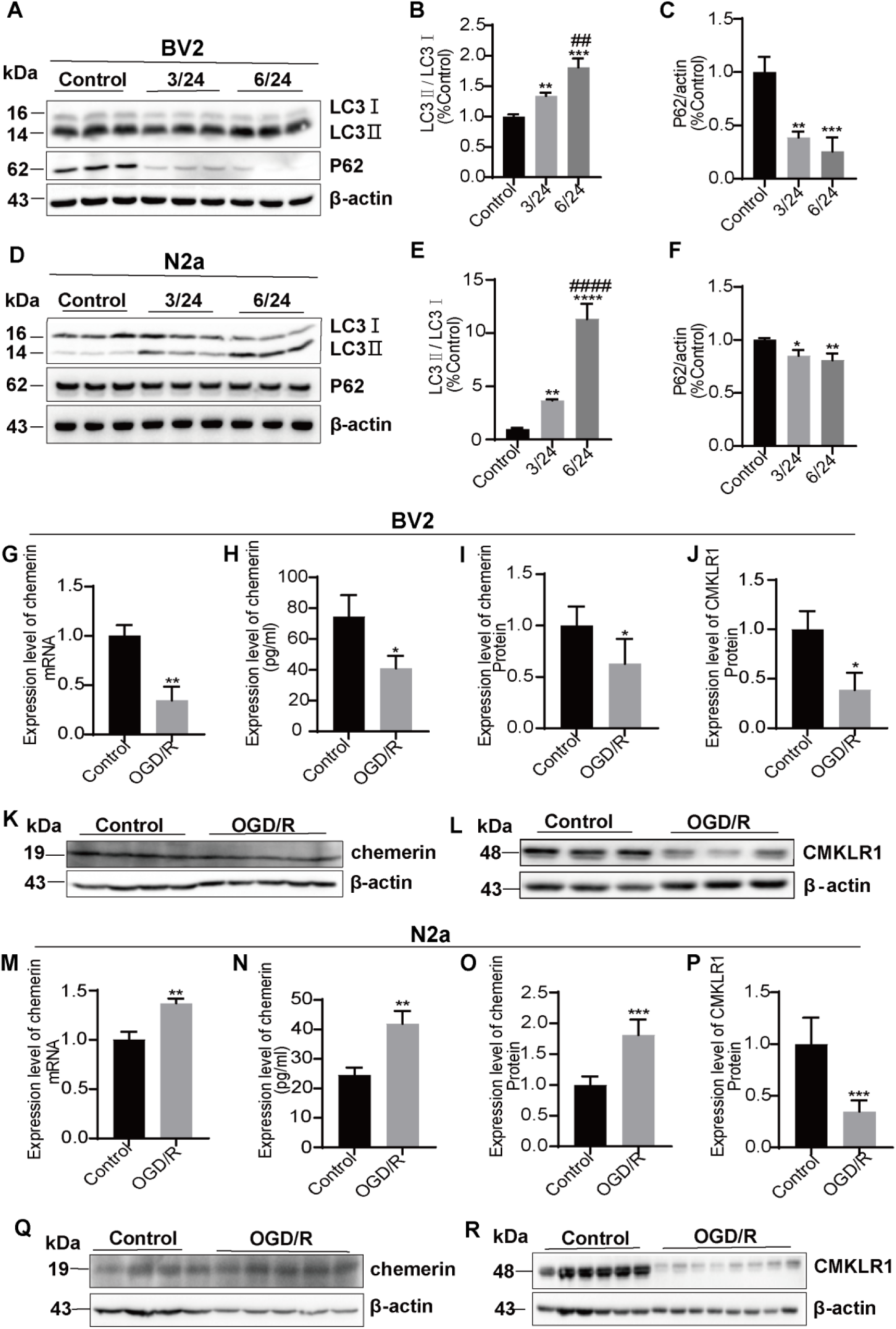
OGD/R treatment altered the expression of LC3, P62, chemerin and CMKLR1 in BV2 and N2a cells. (A-F) Representative blots and quantification of the protein expression of LC3 and P62 in the control, 3/24 (OGD for 3 h and then reoxygenation culture for 24 h) and 6/24 (OGD for 6 h and then reoxygenation culture for 24 h) groups of BV2 and N2a cells (n=3 respectively). (G, M) The mRNA expression level of chemerin in BV2 cells and in N2a cells (n=3 respectively). (H, N) Chemerin expression levels in the culture medium of BV2 cells and in N2a cells (n=3 respectively). (I-L) Representative blots and quantification of the protein expression of chemerin and CMKLR1 in BV2 cells (n=3-5). (O‒R) Representative blots and quantification of the protein expression of chemerin and CMKLR1 in N2a cells (n=4‒8). \*\*\*\**P*<0.0001, \*\*\**P*<0.001, \*\**P* < 0.01, \**P*<0.05 vs. control group; ###*P*< 0.001, ##*P*<0.01, #*P*<0.05 vs. the 3/24 group. The data are presented as the mean±SD.

We next investigated OGD/R-induced changes (6 h OGD/24 h) in chemerin and CMKLR1 expression in BV2 microglia and N2a cells. In BV2 microglia, OGD/R significantly decreased chemerin expression: mRNA by 65.9% (P=0.0029, Fig. 1G), secreted protein measured by ELISA in the cell culture medium by 45% (P=0.022, Fig. 1H), and cellular protein measured by Western blotting by 37.2% (P=0.04, **Fig. 1I,K**). CMKLR1 protein expression was similarly reduced by 61% (P=0.013, **Fig. 1J,L**) by OGD/R in BV2 microglia. In contrast, N2a cells showed opposite responses to OGD/R: chemerin mRNA increased by 37.1% (P=0.0023, **Fig. 1M**), secreted protein in the cell culture medium by 70% (P=0.004, **Fig. 1N**), and cellular protein by 80% (P=0.0007, **Fig. 1O,Q**). CMKLR1 protein expression in N2a cells decreased by 66% after OGD/R (P=0.0001, **Fig. 1P,R**). These results demonstrate that OGD/R downregulating CMKLR1, P62 levels and LC3II/LC3I ratio in BV2 microglia and N2a neuronal cells, and differentially regulates chemerin expression in BV2 microglia and N2a neuronal cells,

### 3.2 Chemerin treatment exacerbated the OGD/R-induced decrease in mitochondrial fission and increase in fusion in BV2 microglia or N2a cells

We hypothesize that chemerin treatment may influence on the mitochondrial fusion/fission in OGD/R-treated BV2 microglia and N2a cells. Initial viability assays confirmed that recombinant chemerin (10– 120 ng/mL) during reoxygenation did not affect cell survival in either cell type (chemerin+OGD/R vs OGD/R; **SFig. 1A,B**). Thus in the following experiment, recombinant chemerin up to 120 ng/mL was used. We observed that in BV2 microglia, OGD/R (6 h OGD/24 h) significantly decreased Drp1 and Fis1 levels by 55.2% (P<0.0001) and 19.4% (P=0.038), and increased Mfn1 level by 43.5% (P=0.009) compared to control group; while the level of Mfn2 remained unchanged by OGD/R (**Fig. 2A-G**). In N2a cells, OGD/R significantly decreased Drp1 and Fis1 expression by 24.6% (P=0.022) and 66.7% (P=0.0001), respectively, and increased Mfn1 level by 111% (P<0.0001) compared to controls. No significant change was observed in Mfn2 level after OGD/R (**Fig. 2H-M**).

**Fig 2.**
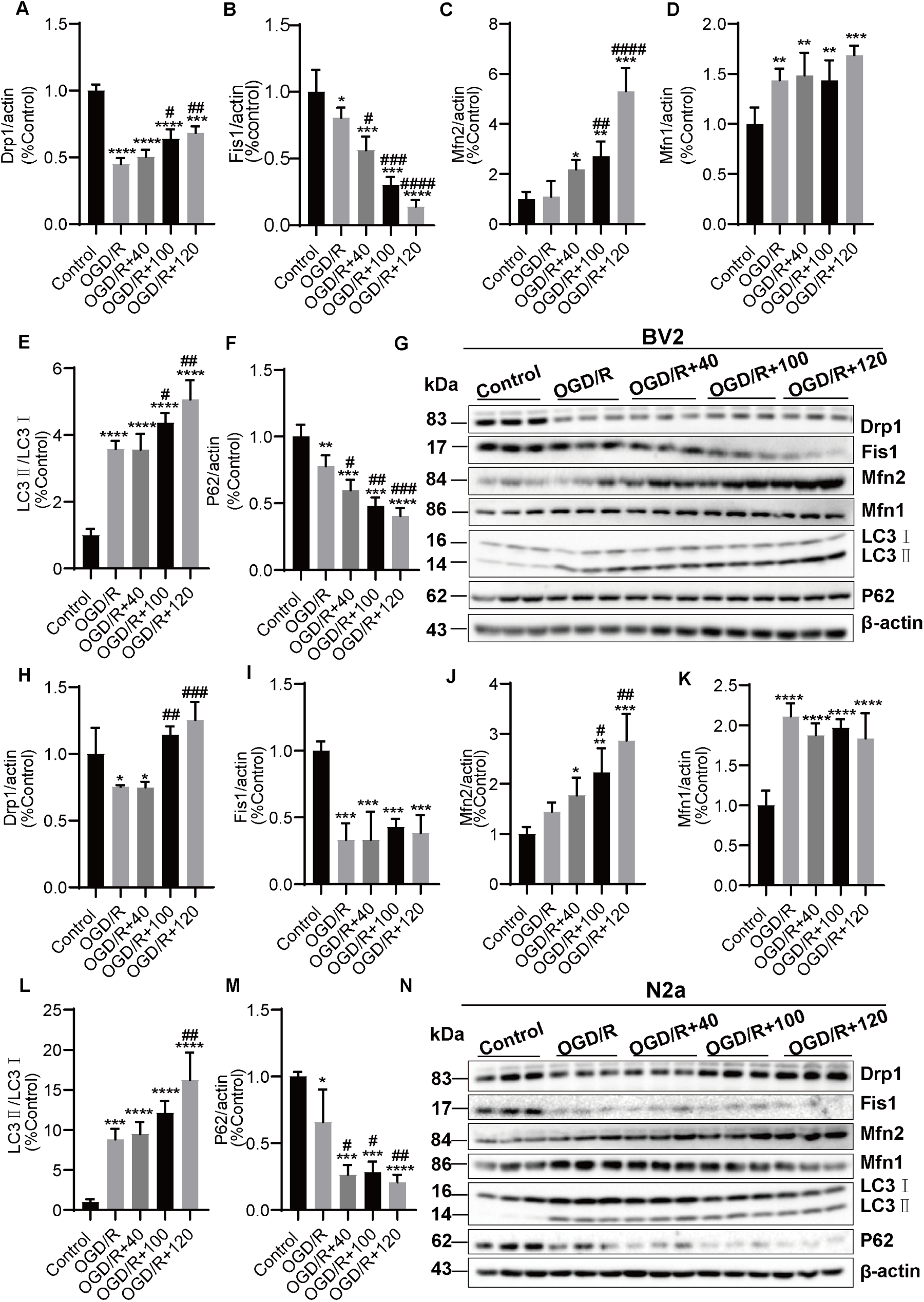
Chemerin exacerbated the OGD/R-induced changes in mitochondrial fusion and fission in BV2 microglia or N2a cells. (A-G) Representative blots and quantification of the protein expression of Drp1, Fis1, Mfn2, Mfn1, LC3 and P62 in the control, OGD/R and OGD/R+chemerin (40, 100 and 120 ng/mL) groups of BV2 cells and (**H-N**) in N2a cells (n=3 respectively). \*\*\*\**p*<0.0001, \*\*\**p*<0.001, \*\**p*<0.01, \**p*<0.05 vs. the control group; ###*p*<0.001, ##*p*<0.01, #*p*<0.05 vs. the OGD/R group. The data are presented as the mean±SD.

We found that in BV2 microglia, treatment with recombinant chemerin dose-dependently modulated these mitochondrial protein levels: Drp1 level was increased by 19% (100 ng/mL, P=0.02) and 23.5% (120 ng/mL, P=0.003) in chemerin+OGD/R vs OGD/R group with no effect at 40 ng/mL (**Fig. 2A,G**). Fis1 level was reduced by 24.4% (40 ng/mL, P=0.013), 50.2% (100 ng/mL, P=0.0001), and 66.5% (120 ng/mL, P<0.0001; **Fig. 2B,G**) in chemerin+OGD/R vs OGD/R group. Mfn2 level was elevated by 161.2% (100 ng/mL, P=0.009) and 420.2% (120 ng/mL, P<0.0001) in chemerin+OGD/R vs OGD/R group, with no effect at 40 ng/mL (Fig. 2C,G). The Mfn1 level was not affected at the three doses of chemerin (**Fig. 2D,G**). Similarly in N2a neuronal cell, chemerin treatment exerted a similar dose-dependent effects on mitochondrial dynamics: Drp1 level was increased by 38.9% (100 ng/mL, P=0.002) and 49.9% (120 ng/mL, P=0.0002) in chemerin+OGD/R versus OGD/R group, with no effect at 40 ng/mL (**Fig. 2H,N**). Mfn2 level was elevated by 78.3% (100 ng/mL, P=0.028) and 142% (120 ng/mL, P=0.001), in chemerin+OGD/R vs OGD/R group with no effect at 40 ng/mL (**Fig. 2J,N**) The levels of Fis1 and Mfn1 remained unchanged at all three tested concentrations of chemerin in N2a cells (40, 100, 120 ng/mL; **Fig. 2I,K,N**).

### 3.3 Chemerin dose-dependently promoted autophagy in OGD/R-treated BV2 microglia and N2a cells

Next we assessed the effect of chemerin treatment on the OGD/R related autography. We found that in BV2 microglia, OGD/R significantly increased LC3II/LC3I ratio by 258% (P<0.0001), and reduced P62 level by 22.1% (P=0.005) versus controls. Chemerin treatment further modulated these effects in a dose-dependent manner: the LC3II/LC3I ratio was increased by 78% (100 ng/mL, P=0.032) and 149% (120 ng/mL, P=0.001), with no effect at 40 ng/mL (**Fig. 2E,G**) compared with the OGD/R alone group. The level of P62 was decreased by 18.1% (40 ng/mL, P=0.015), 29.6% (100 ng/mL, P=0.001), and 37.2% (120 ng/mL, P=0.00013) respectively compared with the OGD/R alone group (**Fig. 2F,G**)

In N2a cells, more pronounced baseline responses to OGD/R was observed: LC3II/LC3I ratio was increased by 782% (P=0.0005) and P62 was decreased by 34% (P=0.04). The effect of chemerin on neuron differed from that in BV2 microglia. For LC3II/LC3I ratio, only the highest dose (120 ng/mL) of chemerin showed significant elevation (738%, P=0.001), while 40, or 100 ng/mL chemerin were ineffective (**Fig. 2L-N**). While all doses of chemerin reduced expression of P62 compared with the OGD/R group (40 ng/mL: 39.4%, P=0.01; 100 ng/mL: 37.4%, P=0.02; 120 ng/mL: 45%, P=0.007) (**Fig. 2L-N**).

### 3.4 CMKLR1 increased mitochondrial fusion and reduced fission in OGD/R-treated BV2 microglia and N2a cells

Next, we assessed how CMKLR1 affects mitochondrial fusion and fission under OGD/R conditions in BV2 microglia and in N2a cells. In BV2 microglia: the levels of Drp1 (*P*=0.016, *P*=0.015, *P*=0.001, **Fig. 3B, F**) and Fis1 (*P*=0.002, *P*=0.001, *P*<0.0001, **Fig. 3C, F**) were lower in the CMKLR1, vector+OGD/R, CMKLR1+OGD/R groups vs vector group. The Fis1 level was lower in the CMKLR1+OGD/R group than in CMKLR1, and vector+OGD/R groups (*P*=0.019, *P*=0.026, **Fig. 3C, F**). The levels of Mfn2 (*P*=0.003, *P*=0.0005, *P*=0.000002) and Mfn1 (*P*=0.014, *P*=0.002, *P*=0.00004) were greater in the CMKLR1, vector+OGD/R, CMKLR1+OGD/R groups vs vector group (**Fig. 3D-F**). Mfn2 (*P*=0.00059, *P*=0.005) and Mfn1 (*P*=0.001, *P*=0.005) levels were greater in the CMKLR1+OGD/R group compared with the CMKLR1 and vector+OGD/R groups (**Fig. 3D-F**).

**Fig. 3.**
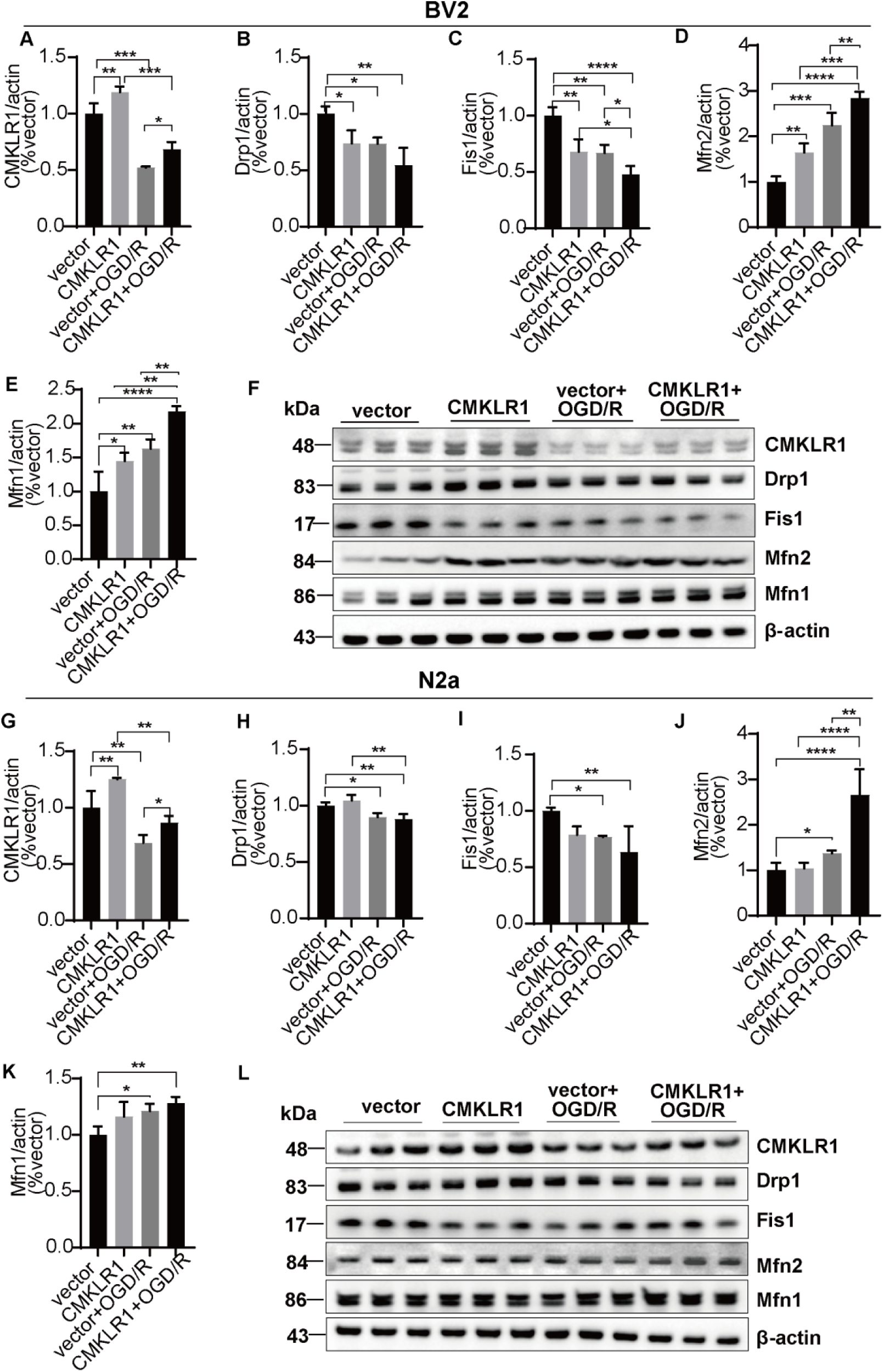
CMKLR1 promoted mitochondrial fusion and reduced fission in OGD/R-treated BV2 microglia and N2a cells. (A, **G)** Representative blots and quantification of the protein expression of CMKLR1 in BV2 microglia and N2a cells (n=3 respectively). **(B-F)** Representative blots and quantification of the protein expression of Drp1, Fis1, Mfn2 and Mfn1 in BV2 cells and **(H-L)** in N2a cells (n=3 respectively). \*\*\*\**p*<0.0001, \*\*\**p*<0.001, \*\**p* < 0.01, \**p*<0.05. The data are presented as the mean±SD.

In contrast, in N2a cells, the levels of Drp1 were lower in vector+OGD/R and CMKLR1+OGD/R groups compared to the vector group (*P*=0.016, *P*=0.007, **Fig. 3H, L**). the level of Fis1 was lower in the vector+OGD/R and CMKLR1+OGD/R groups compared with vector group (*P*=0.048, *P*=0.006, **Fig. 3I, L**), and not different from the CMKLR1 group (*P*=0.066). The level of Mfn2 was greater in the vector+OGD/R and CMKLR1+OGD/R groups vs vector group (*P*=0.034, *P*=0.000062, **Fig. 3J, L**). The Mfn1 level was greater in vector+OGD/R and CMKLR1+OGD/R groups vs vector group (*P*=0.018, *P*=0.004, **Fig. 3K, L**), and not for CMKLR1 group (*P*=0.052). Mfn2 level (but not Mfn1) was greater in the CMKLR1+OGD/R group (*P*=0.00008, *P*=0.001) compared with CMKLR1 and vector+OGD/R groups (**Fig. 3J-L**). These findings indicate that both CMKLR1 overexpression and OGD/R synergically decreased mitochondrial fission and promoted mitochondrial fusion in BV2 microglia and N2a cells, with CMKLR1 showed stronger effect on microglia than neurons.

### 3.5 CMKLR1 increased autophagy in OGD/R-treated BV2 microglia and N2a cells, and mitophagy in N2a cells

We next examined CMKLR1’s role in regulating mitophagy under OGD/R conditions. Quantitative analysis revealed distinct patterns in microglia versus neuronal cells: In BV2 microglia, LC3II/LC3I ratio was elevated in CMKLR1 (P=0.00013), vector+OGD/R (P=0.001), and CMKLR1+OGD/R (P<0.0001) groups vs vector controls (**Fig. 4A,E**). The CMKLR1+OGD/R group showed highest LC3II/LC3I levels (P=0.049 vs CMKLR1; P=0.002 vs vector+OGD/R). Similarly, Pink1 was greater in CMKLR1, vector+OGD/R and CMKLR1+OGD/R groups (P=0.006, P=0.001, P=0.00021) vs vector group. The level of Pink1 was greater in CMKLR1+OGD/R group vs CMKLR1 group (P=0.029, **Fig. 4C, E**). In addition, OGD/R reduced P62 (P=0.002) and increased Parkin (P≤0.013) versus controls, while CMKLR1 overexpression showed no significant effect on P62 or Parkin (**Fig. 4B,D,E**) in BV2 microglia.

**Fig. 4.**
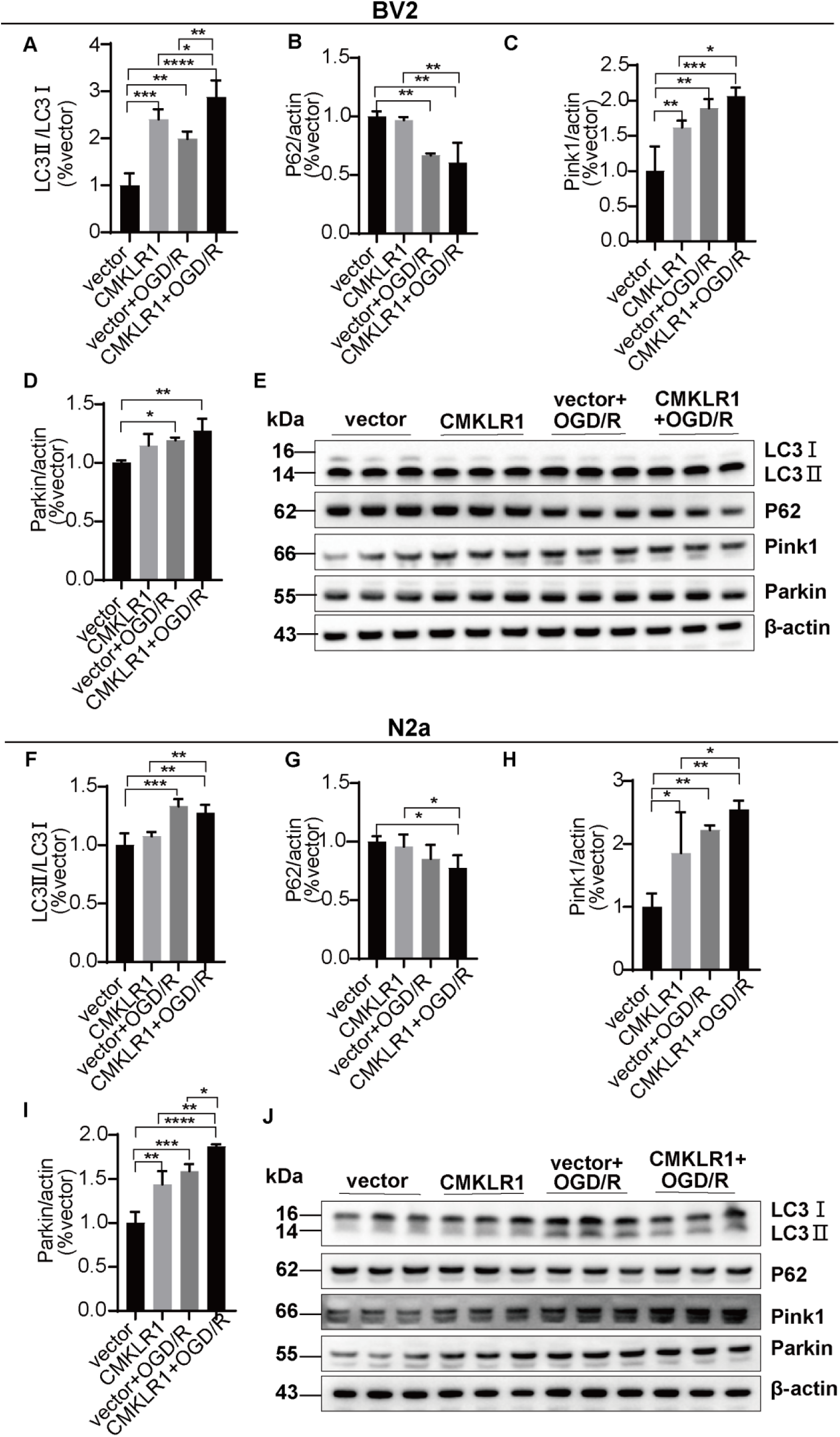
CMKLR1 increased mitophagy levels in OGD/R-treated BV2 microglia and N2a cells. (A-E) Representative blots and quantification of the protein expression of LC3, P62, Pink1 and Parkin in BV2 cells (n=3). **(F-J)** Representative blots and quantification of the protein expression of LC3, P62, Pink1 and Parkin in N2a cells (n=3). \*\*\*\**p*<0.0001, \*\*\**p*<0.001, \*\**p* < 0.01, \**p*<0.05. The data are presented as the mean±SD.

In N2a cell, the level of LC3II/LC3I (*P*=0.0004 and *P*=0.001) was greater in vector+OGD/R and CMKLR1+OGD/R groups vs vector group. Similarly, the levels of Pink1 (*P*=0.017, *P*=0.003 and *P*=0.001) and Parkin (*P*=0.001, *P*=0.0001 and *P*<0.0001) were greater in the CMKLR1, vector+OGD/R and CMKLR1+OGD/R groups vs vector group. The levels of LC3II/LC3I, and Pink1 were greater in the CMKLR1+OGD/R group than CMKLR1 group in N2a cells (*P*=0.002, *P*=0.043, **Fig. 4F, H, J**). Unlike in microglia, Parkin level was greater than in CMKLR1+OGD/R group compared to CMKLR1 and vector+OGD/R groups in N2A cells (*P*=0.001, *P*=0.011, **Fig. 4I, J**); P62 protein level did not change in different conditions in N2a cells (**Fig. 4G, J**). These findings indicated that CMKLR1 activation and OGD/R synergistically enhance autophagy in both cell types, as evidenced by increased LC3II/LC3I and Pink1 levels. In addition, we observed cell-type-specific differences in Parkin, P62 regulation, suggesting distinct mitophagy activation mechanisms between microglia and neurons.

### 3.6 Chemerin reduced OGD/R-induced oxidative stress in BV2 microglia and N2a cells

We investigated chemerin’s effects on oxidative stress markers using flow cytometry and immunofluorescence to detect changes in the levels of ROS, 4-HNE and 8-OHdG. Our analysis revealed both shared and cell-type-specific protective effects. In BV2 microglia, OGD/R increased the level of ROS by 59.6% (*P*<0.0001) compared with the control. Chemerin at 120 ng/mL (not at 40 or 100 ng/mL), decreased the level of ROS in the OGD/R group by approximately 11.7% (*P*=0.038, **Fig. 5A, D**). OGD/R increased the expression of the oxidative stress markers 4-HNE and 8-OHdG by 312% (*P*<0.0001) and 303% *(P*<0.0001) compared with the control. Chemerin reduced 4-HNE (at 100, and 120 ng/mL, *P*=0.001, *P*=0.0001) and 8-OHdG expression (and at 40, 100, and 120 ng/mL, *P*=0.00012, *P*<0.0001, *P*<0.0001) in OGD/R-treated BV2 microglia (**Fig. 5B, C, E, F**).

**Fig. 5.**
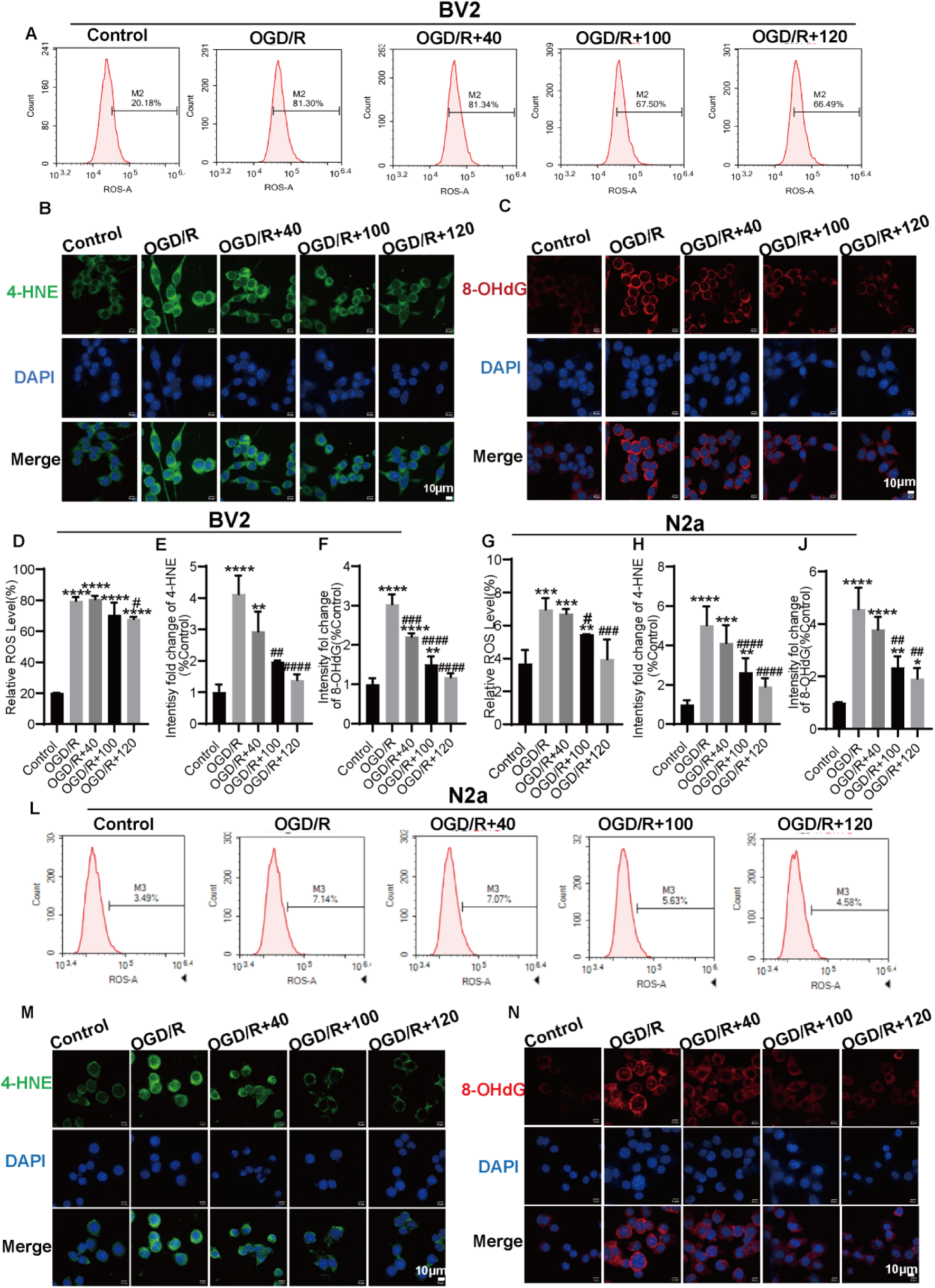
Chemerin decreased oxidative stress in OGD/R-treated BV2 microglia and N2a cells. (A, **L)** Peak plot of ROS in BV2 and N2a cells via flow cytometry. **(B, M)** Fluorescence staining of 4-HNE (green) in the control, OGD/R, and OGD/R+chemerin (40, 100, 120 ng/mL) groups of BV2 and N2a cells (n=3 respectively). Nuclei are stained with DAPI (blue), scale bar=10 μm. (**D, G)** Analysis of the ratio of ROS-positive cells to total cells. **(E, H)** Mean fluorescence intensity analysis of 4-HNE in BV2 and N2a cells. **(C, N)** Fluorescence staining of 8-OHdG (red) in BV2 and N2a cells (n=3 respectively). Nuclei were stained with DAPI (blue), scale bar=10 μm. **(F, J)** Mean fluorescence intensity analysis of 8-OHdG in BV2 and N2a cells. \*\*\*\**P*<0.0001, \*\*\**p*<0.001, \*\**p*<0.01, \**p*<0.05 vs the control group; ####*P*<0.0001, ###*p* < 0.001, ##*P*<0.01, #*P*<0.05 vs the OGD/R group. Data are presented as the mean ± SD.

In N2a cells. we found that OGD/R increased the level of ROS by 327% compared with control group (*P*=0.0002). Chemerin at 100 and 120 ng/mL (not 40 ng/mL), decreased the level of ROS in OGD/R group by 151% (*P*=0.03) and 301% (*P*=0.0004), respectively (**Fig. 5G, L**)., OGD/R increased the expression of 4-HNE and 8-OHdG by 402% (*P*<0.0001) and 356% (*P*<0.0001) compared with control. Similarly, chemerin (at 100, and 120 ng/mL, but not at 40 ng/mL) reduced 4-HNE (*P*<0.0001, *P*<0.0001) and 8-OHdG expression (*P*=0.004 and *P*=0.001) in OGD/R-treated N2a cells (**Fig. 5G-N**).

### 3.7 CMKLR1 overexpression increased oxidative stress in BV2 microglia and N2a cells

Next we evaluated the effect of CMKLR1 overexpression on oxidative stress by using MitoSOX Red and 8-OHdG assays. MitoSOX Red is a fluorescent probe that specifically targets the mitochondria of living cells with membrane permeability and can rapidly and specifically target mitochondria for the selective detection of superoxides in mitochondria. We found that MitoSOX levels in the BV2 microglia were greater in the CMKLR1, vector+OGD/R and CMKLR1+OGD/R groups compared with vector group (*P*=0.00011, *P*<0.0001 and *P*<0.0001, respectively). CMKLR1+OGD/R group showed high MitoSOX expression in BV2 microglia compared with CMKLR1 and vector+OGD/R groups (*P*<0.0001, *P*<0.0001, **Fig. 6A, B**). The expression of 8-OHdG in BV2 microglia was greater in the vector+OGD/R and CMKLR1+OGD/R groups vs vector group (*P*=0.047 and *P*=0.00049). 8-OHdG levels in BV2 microglia were significantly greater in the CMKLR1+OGD/R group compared with CMKLR1 and vector+OGD/R groups (*P*=0.0003, *P*=0.014) (**Fig. 6E, F**).

**Fig 6.**
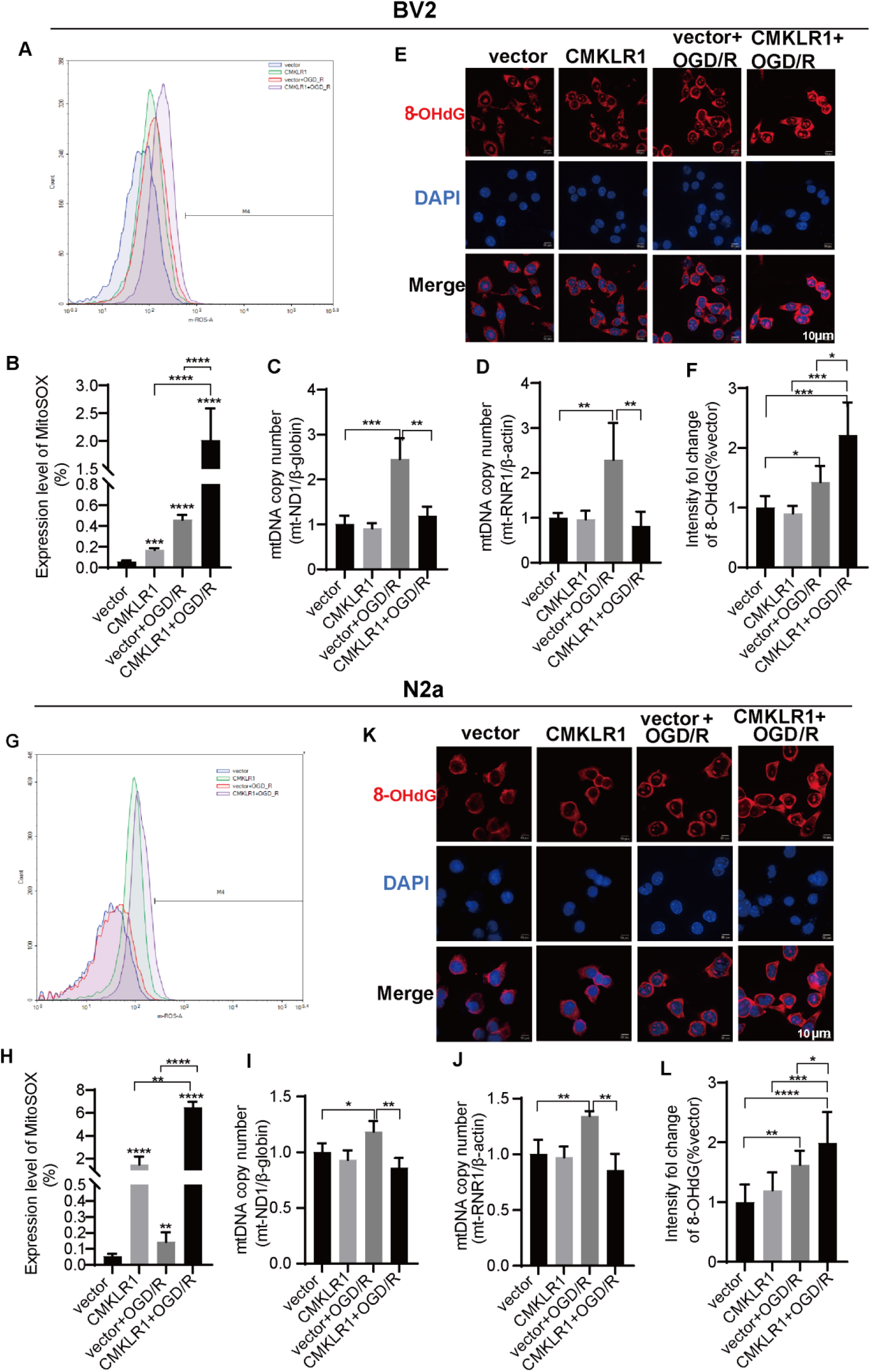
CMKLR1 overexpression increased oxidative stress and decreased mtDNA copy number in OGD/R-treated BV2 microglia and N2a cells. (A, B, G, **H)** Representative flow cytometry images and quantitative statistics of the percentage of MitoSOX-positive BV2 and N2a cells (n=3 respectively). **(C, D, I, J)** Changes in mtDNA copy number in BV2 and N2a cells (n=3 respectively). **(E, K)** Fluorescence staining of 8-OHdG (red) in BV2 and N2a cells. Nuclei are stained with DAPI (blue) (n=3 respectively), scale bar=10 μm. **(F, L)** Mean fluorescence intensity analysis of 8-OHdG in BV2 and N2a cells. \*\*\*\**p*<0.0001, \*\*\**p*<0.001, \*\**p* < 0.01, \**p*<0.05. The data are presented as the mean±SD.

In N2a cells, overexpression of CMKLR1 also increased the expression of MitoSOX. The levels of MitoSOX were greater in the CMKLR1, vector+OGD/R and CMKLR1+OGD/R groups (*P*<0.0001, *P*=0.008, *P*<0.0001, respectively) than vector group. CMKLR1+OGD/R group showed high MitoSOX expression compared to CMKLR1 group and vector+OGD/R groups (*P*=0.001, *P*<0.0001, **Fig. 6G, H**). The expression of 8-OHdG was greater in the vector+OGD/R and CMKLR1+OGD/R groups vs vector group (*P*=0.001, *P*<0.0001), and greater in the CMKLR1+OGD/R group compared to CMKLR1 group and vector+OGD/R groups (*P*=0.00011 and *P*=0.048) (**Fig. 6K, L**).

### 3.8 CMKLR1 overexpression decreased mtDNA copy number, led to mitochondrial respiratory disturbances in BV2 microglia and N2a cells

Next we evaluated the effect of CMKLR1 overexpression on mitochondrial biogenesis and mitochondrial respiration. In BV2 microglia, the mtDNA copy numbers (mt-ND1/β-globin and mt-RNR1/β-actin) in the vector+OGD/R group were greater in BV2 microglia (*P*=0.00021, *P*=0.009 vs vector group). CMKLR1 overexpression reduced the mtDNA copy number (mt-ND1 and mt-RNR1) under OGD/R conditions in BV2 microglia (CMKLR1 +OGD vs vector+OGD/R group, *P*=0.001 and *P*=0.004) (**Fig. 6C, D**). No difference was observed in basal respiration, proton leakage, maximal respiration, spare respiratory capacity, nonmitochondrial oxygen consumption or ATP-related respiration in the CMKLR1 group compared with vector group in BV2 microglia. In BV2 microglia, OGD/R reduced basal respiration, maximal respiration, spare respiratory capacity and ATP-related respiration (*P*=0.047, *P*=0.007, *P*=0.001, *P*=0.025, vector+OGD/R vs vector group). CMKLR1 overexpression led to higher nonmitochondria consumption in BV2 microglia (*P*=0.033, CMKLR1+OGD/R vs vector+OGD/R group, **Fig. 7A-D**).

**Fig. 7.**
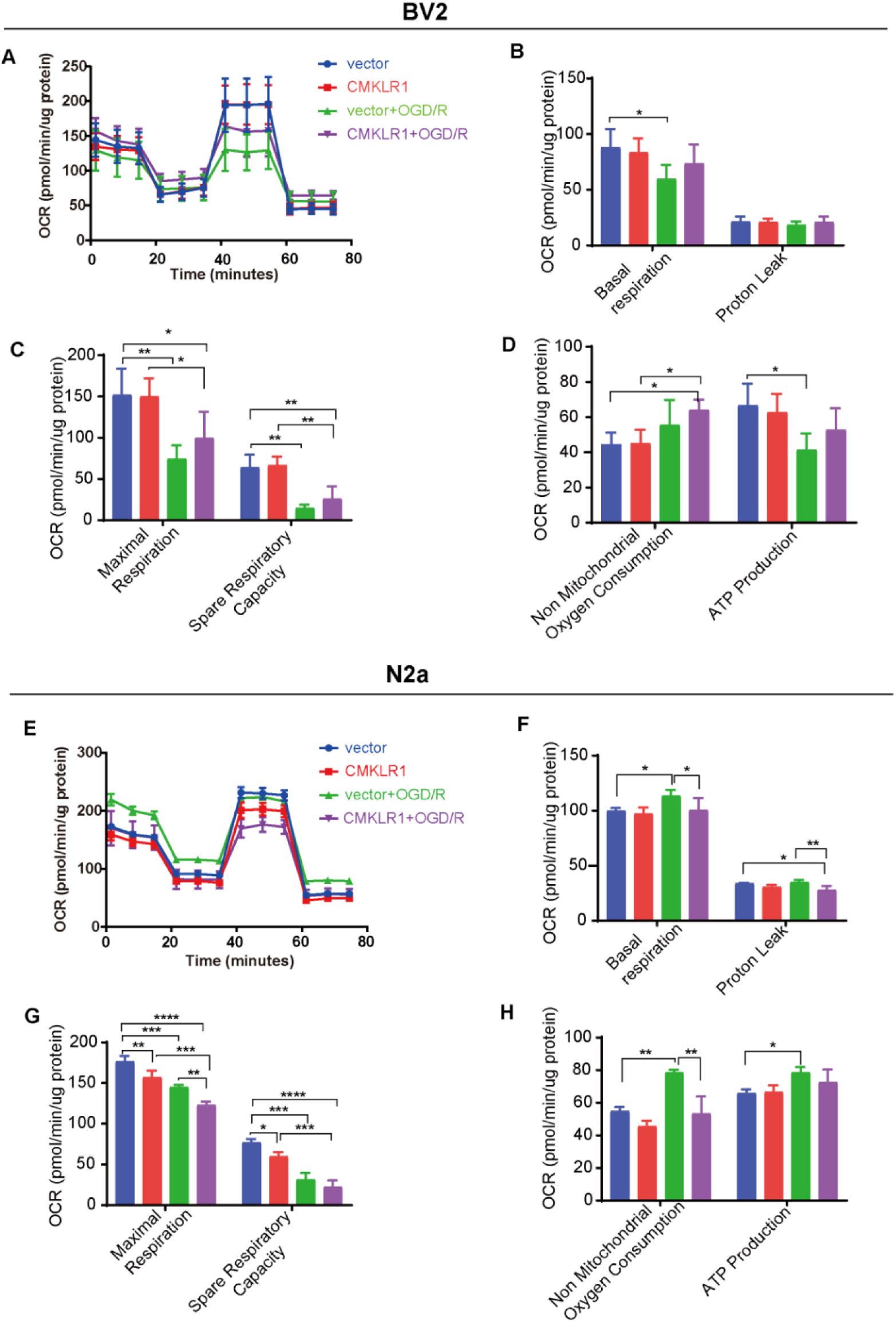
CMKLR1 overexpression led to mitochondrial respiratory disturbances in OGD/R-treated BV2 and N2a cells. (A, E) Mitochondrial stress detection via real-time images of BV2 and N2a cells. (B-D, F-H) Statistical analysis of key parameters of mitochondrial pressure (n=3 respectively). \**P*<0.05, \*\**P*<0.01, \*\*\*\**P*<0.001, \*\*\*\**P*<0.0001. The data are presented as the mean± SD.

In N2a cells, the mtDNA copy numbers (mt-ND1/β-globin and mt-RNR1/β-actin) were increased after OGD/R (*P*=0.032 and *P*=0.005, vs vector group), which is reversed by CMKLR1 overexpression (*P*=0.002, *P*=0.001, CMKLR1+OGD vs vector+OGD/R group, **Fig. 6I, J**). CMKLR1 group and vector group did not differ in the basal respiration, proton leakage, or ATP-related respiration in N2a cells. The maximum respiration and spare respiration capacity were significantly lower in the CMKLR1 group than vector group (*P*=0.003 and *P*=0.014). OGD/R increased basal respiration (*P*=0.038), nonmitochondrial oxygen consumption (*P*=0.001) and ATP-related respiration (*P*=0.012), whereas reduced the maximal and spare respiratory capacities (*P*=0.00016, *P*=0.00032, vector+OGD/R vs vector group) in N2a cells. CMKLR1 overexpression reduced basal respiration (*P*=0.047), proton drain (*P*=0.006), maximal respiration (*P*=0.002), nonmitochondrial oxygen consumption (*P*=0.001, CMKLR1+OGD/R group vs vector+OGD/R group, **Fig. 7E-H**), but not spare respiratory capacity or ATP-related respiration. Taken together, these findings indicate that CMKLR1 differentially affect the mitochondrial respiratory function after OGD/R in BV2 microglia and N2a neuronal cells.

### 3.9 Opposing effects of chemerin and CMKLR1 on OGD/R-induced apoptosis in BV2 microglia and N2a neuronal cells

We measured the effects of OGD/R on the apoptosis of BV2 and N2a cells. The percentages of annexin V-FITC-and PI-positive cells were analyzed using flow cytometry. FITC-positive cells indicate early apoptotic cells, whereas FITC-and PI-positive cells indicate late apoptotic cells. The percentage of apoptotic cells is equal to the percentage of early apoptotic cells plus late apoptotic cells. The apoptosis rate was increased by approximately 12.3% (*P<*0.0001) in the OGD/R group of BV2 microglia and 3.25% (*P*=0.002) in the N2a cell group vs control group. In addition, 120 ng/mL chemerin significantly reduced the degree of apoptosis caused by OGD/R in BV2 microglia (P=0.012) and N2a cells (P=0.015) (**Fig. 8A-D**).

**Fig 8.**
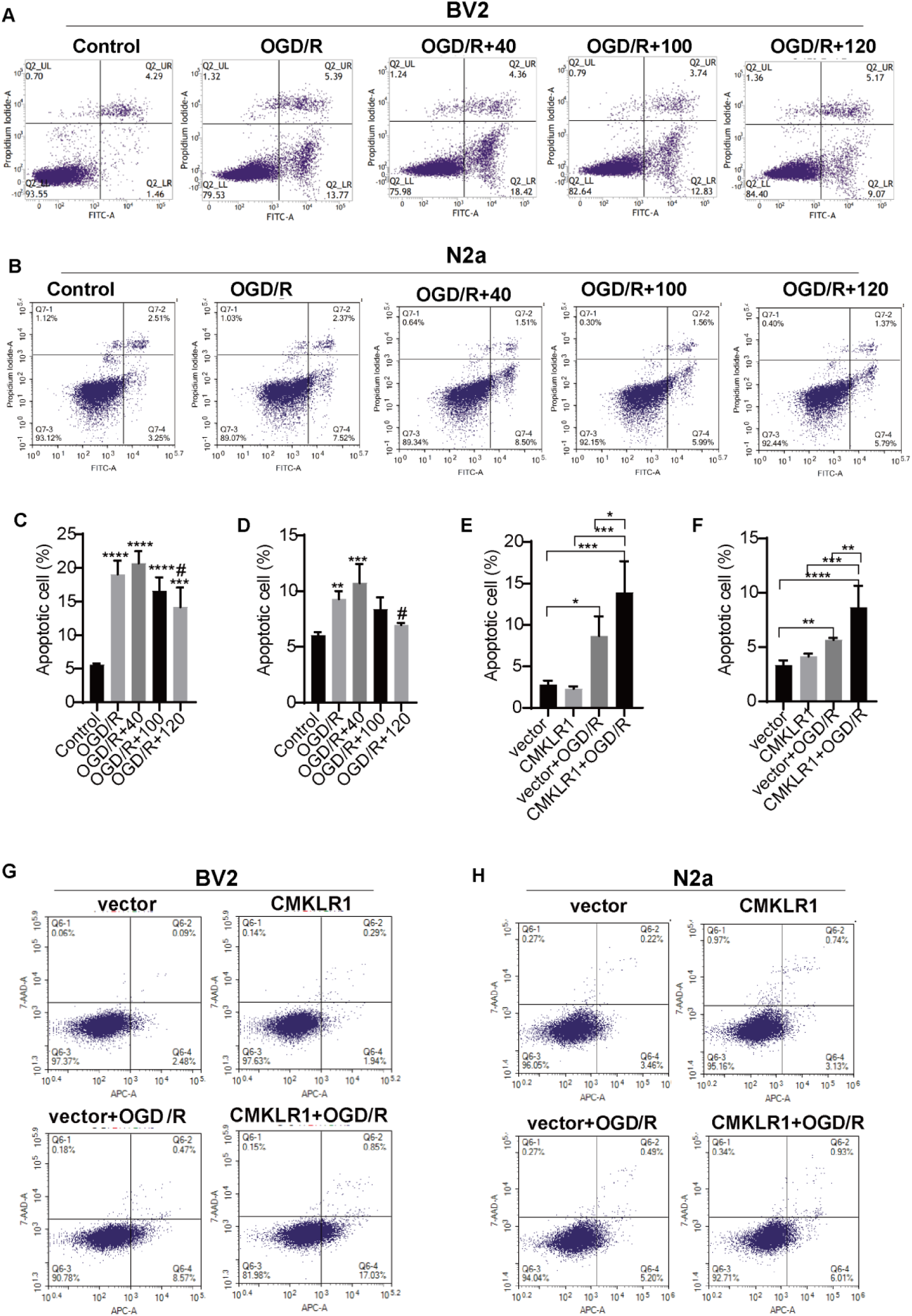
Differential effects of chemerin and CMKLR1 on OGD/R-induced apoptosis in BV2 microglia and N2a cells. (A, B) Representative flow cytometry images of BV2 and N2a apoptosis. (C, D) Quantitative statistics of the percentage of apoptotic BV2 cells and N2a cells in the control, OGD/R and OGD/R+chemerin (40, 100 and 120 ng/mL) groups (n=3 respectively); \*\*\**P*<0.001, \*\**P*<0.01 vs the control group; #*P*<0.05 vs the OGD/R group. The data are presented as the means ± SDs. (E-F, G-H) Representative flow cytometry images of apoptosis and quantitative statistics of the percentages of apoptotic BV2 and N2a cells in the vector, CMKLR1, vector+OGD/R and CMKLR1+OGD/R groups, n=3, \**P*<0.05, \*\**P*<0.01, \*\*\*\**P*<0.001, \*\*\*\**P*<0.0001. The data are presented as the mean± SD.

Next, we assessed the influence of CMKLR1 overexpression and OGD/R on the apoptosis of BV2 and N2a cells. Annexin V-APC and 7-AAD-labeled apoptosis kits were used, and total apoptosis rates were detected using flow cytometry. The percentage of apoptotic BV2 microglia and N2a cells in the CMKLR1 group did not differ from that in the vector group. The degree of apoptosis in BV2 microglia was significantly greater in the vector+OGD/R and CMKLR1+OGD/R groups than in the vector group (*P*=0.013 and *P*=0.00031). CMKLR1 and OGD/R synergistically increased apoptosis in BV2 microglia (*P*=0.000236 CMKLR1 vs CMKLR1+OGD/R group; *P*=0.022 vector+OGD/R vs CMKLR1+OGD/R group; **Fig. 8E, G**) The degree of apoptosis was significantly greater in the vector+OGD/R and CMKLR1+OGD/R groups than in the vector group in N2a cells (*P*=0.005 and *P*=0.00007). CMKLR1 and OGD/R synergistically increased apoptosis in N2a cells (*P*=0.0003 CMKLR1 vs CMKLR1+OGD/R group, *P*=0.006 vector+OGD/R vs CMKLR1+OGD/R group; **Fig. 8F, H**). CMKLR1 and OGD/R synergistically further increased apoptosis. These findings indicated that OGD/R induces apoptosis in both cell types, with microglia showing greater susceptibility. In addition, Chemerin exerts consistent anti-apoptotic effects, and suggesting that cell-type-specific apoptotic pathways are differentially regulated.

## DISCUSSION

We found that OGD/R had opposing cell-type-specific regulation of chemerin expression-decreased in BV2 microglia while increased in N2a neuronal cells, yet consistently reduced CMKLR1 expression and increased autophagy in both cell types. Recombinant chemerin during the reoxygenation phase reduced OGD/R-induced mitochondrial fusion, bioenergetic dysfunction, oxidative stress and apoptosis and increased mitochondrial fission and autophagy in both BV2 microglia and N2a cells. Moreover, CMKLR1 overexpression and OGD/R, showed detrimental effects on BV2 microglia and N2a cells including decreased mitochondrial fission and promoted mitochondrial fusion, mitophagy, and apoptosis, with cell-type-specific differences in Parkin, P62 regulation, suggesting distinct mitophagy activation mechanisms. These findings highlight the complex, cell-specific roles of the chemerin-CMKLR1 system in ischemic injury while demonstrating chemerin’s potential as a multi-target therapeutic agent.

One of the main findings in our study were the cell type-specific change in the levels of chemerin in BV2 microglia and N2a neuroblastoma cells induced by OGD/R. Extensive research has been dedicated to unraveling the mechanisms [42] and therapeutic strategies associated with I/R injury reperfusion therapies [7]. To date, only a few studies have investigated chemerin and CMKLR1 as targets for ischemic stroke. An earlier study has reported increase in ChemR23 after 3 h OGD in using a coculture system [35], which has a different design compared to the 6 h OGD/24 hour reperfusion condition in BV2 microglia in the current study. Microglia play important roles in I/R injury. The postischemic inflammatory response begins within several minutes after the onset of brain ischemia. Secondary neuroinflammation promotes further injury, has a great impact on the outcome of stroke-related mortality and is an important predisposing factor for I/R injury. Several key pathways, including the Notch, phosphatidylinositol 3-kinase/protein kinase B, transforming growth factor-β, nuclear factor-kappa, and Wnt pathways, have emerged as crucial players in I/R injury [42]. An earlier study characterizing the dynamics of the M1/M2 status of BV2 microglia after I/R indicated that at 24 hours, there was an increase in M2 microglia after OGD/R [35]. In addition, neutrophil mobilization triggers changes in microglial function to exacerbate cerebral I/R injury [43]. Significant differences in microvascular density, flow and hypoxia in the ipsi- and contralateral brain hemispheres [44, 45] and the spinal cord of tMCAO model mice have been reported [46] in a mouse model of ischemic stroke [47, 48].Moreover, one fragment of chemerin, the chemerin 15 peptide, has been shown to reduce neuroinflammation after I/R injury [35], inhibit neutrophil-mediated vascular inflammation through ChemR23 [49], mitigate neointimal hyperplasia and accelerate vascular healing [50].

Moreover, we found that chemerin increased the expression of Drp1, and Mfn2, decreased Fis1, without affecting the Mfn1; whereas CMKLR1 reduced Drp1 and Fis1, and increased Mfn1 and Mfn2 in BV2 microglia and N2a cells. Mitochondria are dynamic organelles that maintain the homeostasis of cells through continuous fission, fusion, autophagy, generation, and other key processes that maintain normal mitochondrial function [53]. Mitochondrial dynamics disorders are important features of I/R injury, with the inhibition of mitochondrial fission as a therapeutic target [51]. Mitochondrial fusion is regulated by optic atrophy 1, Mfn1 and Mfn2, whereas mitochondrial fission is regulated mainly by Drp1 and Fis1. However, Mfn2, not Mfn1, is necessary for mitochondrial respiration to adapt to stress conditions, as demonstrated in an earlier study using Mfn2 and Mfn1 conditional knockout mouse models [54]. Chemerin has been reported to increase the expression of Drp1 and Fis1, reduce the expression of the Mfn2 and optic atrophy 1 proteins, and increase mitophagy [29]. In the early stages after ischemia, Drp1-dependent mitophagy has been shown to help clear damaged mitochondria [52]. Increased expression of Drp1 by chemerin treatment may thus increase the clearance of damaged mitochondria. An earlier study shown that defects in Mfn2 caused by CMT2A disease mutation can be rescued by Mfn1, which indicates partial functional redundancy between Mfn1 and Mfn2 [55].

Mitochondria are the main sites of aerobic oxidation and energy conversion in cells [56]. Mitochondrial damage has been shown to be one of the main causes of ischemic neuronal death. Oxidative stress has been shown to mediate chemerin-induced autophagy in endothelial cells [57]. Both dysfunctional mitochondria and ROS produced by mitochondrial oxidative stress can induce mitophagy, and damaged mitochondria regulate mitochondria-dependent apoptosis in ischemic neurons through mitophagy [58]. Here we found that chemerin treatment reduced oxidative stress (ROS, 4-HNE, and 8-OHdG) and apoptosis after OGD/R. Under baseline conditions, CMKLR1-overexpressing BV2 microglia and N2a cells maintained mitochondrial homeostasis, effectively clearing ROS without altering 8-OHdG, mtDNA copy number, or apoptosis. OGD/R disrupted this balance, and increased the levels of mitochondrial ROS, 8-OHdG, and apoptosis. Although the mtDNA copy number initially increased as a compensatory response, CMKLR1 overexpression under OGD/R exacerbated ROS production, overwhelming mitochondrial defenses, respiratory dysfunction, reducing maximal respiration and spare capacity, similarly in BV2 microglia and N2a cells. This resulted in mtDNA damage, reduced copy numbers, and further apoptosis. Furthermore, we showed that mitochondrial respiration was differentially affected in BV2 and N2a cells. In BV2 microglia, OGD/R impaired mitochondrial function, reducing basal/maximal respiration, spare capacity, and ATP production while increasing nonmitochondrial oxygen consumption; whereas in N2a cells, OGD/R alone increased basal respiration, ATP-linked respiration, and nonmitochondrial consumption, suggesting increased energy demands. Despite elevated mtDNA copy numbers in N2a cells, the decline in respiratory flexibility indicated compromised metabolic adaptation. Our findings are consistent with reports that mitochondrial DNA levels increase in the first few hours after stroke ischemia, indicating that mitochondrial biogenesis is part of the poststroke repair mechanism during ischemia [59]. Under physiological conditions, intramitochondrial ROS generated primarily in the inner membrane are efficiently cleared. However, excessive stress surpasses this capacity, triggering ROS overproduction, mtDNA oxidative damage, and reduced mtDNA copy number. Mild stress may temporarily increase mtDNA copy number as a compensatory response, whereas severe stress leads to a decrease in mtDNA copy number [60]. In addition, chemerin has been shown to increase sympathetic outflow and blood pressure using glutamate receptor-mediated ROS generation [61]. The increased nonmitochondrial oxygen consumption in both cell types further supports OGD/R-induced cellular stress.

The different responses of BV2 microglia and N2a cells to OGD/R likely reflect their distinct cell type characteristics. Microglia, which respond first to ischemic injury, exhibit greater sensitivity and morphological changes under OGD/R, leading to pronounced mitochondrial damage. In contrast, N2a cells may still be in a compensatory stress phase, explaining their divergent metabolic responses. Overall, OGD/R impairs mitochondrial respiration, and CMKLR1 overexpression exacerbates this dysfunction, as supported by increased MitoSOX, 8-OHdG, and mtDNA copy number changes, which is consistent with increased apoptosis.

Autophagy is essential for the normal function of mitochondria and the maintenance of cellular homeostasis and plays a key role in regulating mitochondrial biogenesis and cell physiology [62]. The protective effect of autophagy during reperfusion may be attributed to the clearance of mitochondria associated with mitophagy and the inhibition of downstream apoptosis [63]. Thus, enhancing mitophagy has been proposed as a therapeutic strategy to protect neurons from ischemic damage [64]. An imbalance in mitophagy may lead to mitochondrial accumulation, increased oxygen consumption, excessive ROS production, and ultimately, cell death pathways [62]. The role of mitophagy in I/R injury remains debated, with inadequate clearance of damaged mitochondria or excessive degradation of functionally intact mitochondria leading to cell death. Studies have shown that promoting mitophagy has a protective effect on cerebral I/R by clearing damaged mitochondria [65, 66], whereas reducing mitophagy improves mitochondrial function [67]. Here we found that the addition of recombinant chemerin after OGD/R promoted mitophagy in BV2 microglia and N2a cells, which is consistent with the findings of an earlier study [25]. This leads to a reduced level of oxidative stress and apoptosis caused by OGD/R and a protective effect. In contrast, although CMKLR1 activation and OGD/R synergistically enhance autophagy in both cell types, we observed cell-type-specific differences in Parkin regulation only in neuron, suggesting distinct mitophagy activation mechanisms between microglia and neurons. Chemerin and CMKLR1 increased autophagy but exert divergent effects on oxidative stress and apoptosis, likely due to differences in mitophagy dynamics as observed in the current study. The addition of chemerin promoted autophagy during the reperfusion phase, whereas the overexpression of CMKLR1 promoted autophagy before OGD. Thus, the different effects on oxidative stress and apoptosis may also be related to the different time points and pathophysiological microenvironments in which autophagy occurs [63] and require further investigation.

There are several limitations in this study. First, we studied microglia and neurons separately. A mixture of neurons and microglia with single-cell analysis might better mimic the real situation [35]. Second, we examined the effect of chemerin under reoxygenation culture and did not examine the effect under hypoxia culture. Third, we did not further characterize the phenotype of microglia or monitor the changes at different reperfusion time points in the current study [35], as the reperfusion phase is suspected to be the cutoff point when autophagy switches its role in protecting or disrupting neuronal action during stroke. Further comprehensive evaluation at different time points is needed to fully explain the role of chemerin and CMKLR1 in I/R injury.

## CONCLUSION

Our findings demonstrate that OGD/R induces cell-type-specific regulation of the chemerin-CMKLR1 axis, reducing CMKLR1 expression in both microglia and neurons while differentially modulating chemerin expression. Chemerin exerted protective effects against CMKLR1 overexpression that synergized with OGD/R, in mitochondrial dysfunction, oxidative stress, and apoptosis. The complex, cell-specific roles of the chemerin-CMKLR1 system is a potential therapeutic target for I/R injury.

## Declaration

### Author contributions

PYL, NC, XG, and ZWW performed the experiments, and ZZG, XLQ, and ZT helped design the study, analyzed the data, and funded the research. YX designed the study, analyzed the data, and wrote the manuscript. RN revised the manuscript. All the authors read and approved the submission of the final version of the manuscript.

## Funding

This work was supported by the National Natural Science Foundation of China (82260245 to YX, 82360281 to ZT). the Foundation for Guizhou Provincial Science and Technology projects (Nos. ZK [2024] 042 to ZT). Cultivation Foundation of Guizhou Medical University [20NSP069] to YX.

## Conflict of interest

The authors declare no conflicts of interest.

## Supporting information

SFig. 1, STable 1, STable 2, STable 3

## Notes

### Competing Interest Statement

The authors have declared no competing interest.

