## Supplementary material for "Chemerin-CMKLR1 mediated OGD/R induced Mitochondrial Dysfunction, Oxidative Stress, and Autophagy differentially in Microglia and Neurons": SFig. 1, STable 1, STable 2, STable 3

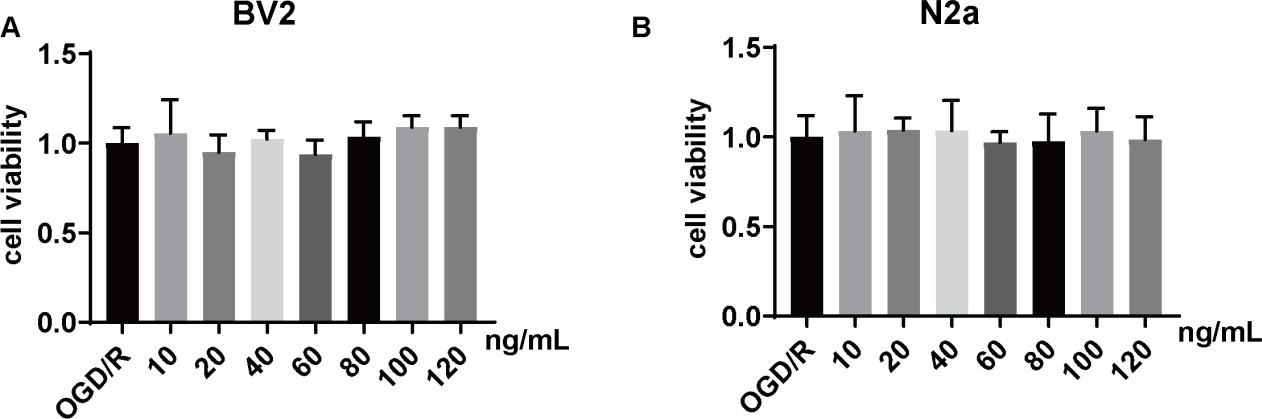

### SFig. 1 Effects of chemerin on the viability of BV2 and N2a cells. (A) Viability of BV2 cells subjected to OGD/R and OGD/R+chemerin protein (10, 20, 40, 60, 80, 100 and 120 ng/mL, n=6 each group). (B) Viability of N2a cells subjected to OGD/R and OGD/R+chemerin protein (10, 20, 40, 60, 80, 100 and 120 ng/mL n=6 each group). The data are presented as the mean±SD.

**STable 1 Antibodies used in this study.**

| Antibody | Host | WB Dilution | IF Dilution | Sources | Catalog No. |
| --- | --- | --- | --- | --- | --- |
| Fis1 | r | 1/1000 |  | GeneTex | GTX111010 |
| Drp1 | r | 1/1000 |  | Abcam | ab184247 |
| Mfn1 | r | 1/1000 |  | Abcam | ab221661 |
| Mfn2 | r | 1/1000 |  | Abcam | ab124773 |
| 4-HNE | r |  | 1:150 | Abcam | ab46545 |
| 8-OHdG | m |  | 1/200 | Abcam | ab62623 |
| SQSTM1/p62 | r | 1/20000 |  | ImmunoWay | YT7058 |
| Pink1 | r | 1/1000 |  | Abcam | ab3707 |
| Parkin | r | 1:20000 |  | ImmunoWay | YT3591 |
| chemerin | r | 1/200 |  | Proteintech | 10216-1-AP |
| CMKLR1 | r | 1:1000 |  | Abcam | ab64881 |
| LC3 | r | 1:2000 |  | Cell Signaling Technology | 12741S |
| β-actin | m | 1:20000 |  | Iffinity | T0022 |
| Goat anti-Rabbit IgG (H+L) Alexa Fluor™ 488 | g |  | 1/500 | Thermo Fisher Scientific | A-11034 |
| Goat anti-Mouse IgG (H+L) Cyanine3 | g |  | 1/500 | Thermo Fisher Scientific | A10521 |
| Anti-mouse IgG, HRP-linked Antibody | Hr | 1/5000 |  | Cell Signaling Technology | 7076S |
| Anti-rabbit IgG, HRP-linked Antibody | g | 1/5000 |  | Cell Signaling Technology | 7074S |

m, mouse; r, rabbit; g, goat; Hr, horse; IF, immunofluorescence; WB, western blot; 4-HNE, 4-hydroxynonenal; 8-OHdG, 8-hydroxy-2’-deoxyguanosine; Fis1, fission 1; Drp1, dynamin-related protein 1; LC3, microtubule-associated protein 1 light chain 3; Mfn1/2, mitofusin 1/2; Pink1, PTEN-induced putative kinase 1; CMKLR1, chemokine-like receptor 1.

| Regents | Sources | Catalog No. |
| --- | --- | --- |
| Mouse recombinant chemerin protein | Sino Biological Inc | 50024-M08H |
| Dulbecco's Modified Eagle Medium | Gibco | 11965092 |
| Sugar-free DMEM | Gibco | 11966025 |
| FBS | Gibco | 10091-148 |
| Bradford Protein Assay Kit | Takara | T9310A |
| Hydrophobic PVDF Transfer Membrane | Merck Millipore Billerica | ISEQ00010 |
| Cell Counting Kit-8 | APE×BIO | K1018 |
| stripping buffer | Solarbio | SW3020 |
| Goat Serum | ZSGB-Bio | ZU-9022 |
| PMSF | Solarbio | P0100 |
| Puromycin | Solarbio | P8230 |
| RIPA buffer (high) | Solarbio | R0010 |
| ROS assay kit | Solarbio | CA1420 |
| Triton X-100 | Solarbio | T8200 |
| Tween-20 | Solarbio | T8220 |
| DAPI | Solarbio | 28718-90-3 |
| TRNzol Universal Reagent | TIANGEN | DP424 |
| TIANamp Genomic DNA Kit | TIANGEN | DP304-02 |
| StarScript Ⅱ RT Mix with gDNA Remover | Genestar | A224-10 |
| 2×RealStar Fast SYBR qPCR Mix | Genestar | A301 |
| MitoSOX^TM^ Red | ThermoFisher | M36008 |
| FITC Annexin V Apoptosis Detection kit | BD | 556547 |
| XF Cell Mito Stress Test Kit | Agilent | 103010-100 |
| Annexin V-APC/7-AAD apoptosis kit | MULTISCIENCES | AP105 |

**STable 2 Reagents used in this study**

PVDF, polyvinylidene fluoride; PMSF, phenylmethylsulfonyl fluoride; RIPA, radio immunoprecipitation assay; ROS, reactive oxygen species; DAPI, 4′,6-diamidino-2-phenylindole.

| gene | Primer sequences (5'→3') |
| --- | --- |
| chemerin | F ATAGTCCACTGCCCAATTCTGAAGC  R TTCGCCAGCCTGTGCTATCTTAATG |
| CMKLR1 | F TCCTGTTCAACATCTTTTTGCC  R CAAGAAGTTGCTGATCTTGCAC |
| mt-ND1 | F GAGCTTTACGAGCCGTAGCC  R CCCGGTTTGTTTCTGCTAGG |
| β-globin | F GACACACAACCCCAGAAACA  R GCCTCACCACCAACTTCATC |
| mt-RNR1 | F AGCAATGAAGTACGCACACA  R TTCCAAGCACACTTTCCAGT |
| β-actin | F TGGAATCCTGTGGCATCCAT  R GCTAGGAGCCAGAGCAGTAA |

**STable 3 Primer sequences**
